# White adipose tissue remodeling in Little Brown Myotis (*Myotis lucifugus*) with white-nose syndrome

**DOI:** 10.1101/2024.06.17.599301

**Authors:** Evan L. Pannkuk, Marianne S. Moore, Shivani Bansal, Kamendra Kumar, Shubhankar Suman, Daryl Howell, Joseph A. Kath, Allen Kurta, DeeAnn M. Reeder, Kenneth A. Field

**Affiliations:** Department of Biochemistry and Molecular & Cellular Biology, Georgetown University Medical Center, Washington, DC 20057, United States of America; Center for Metabolomic Studies, Georgetown University, Washington, DC 20057, United States of America; Department of Oncology, Lombardi Comprehensive Cancer Center, Georgetown University Medical Center, Washington, DC 20057, United States of America; Department of Biological Sciences, University of the Virgin Islands, St. Thomas, USVI 00802; Iowa Department of Natural Resources, Des Moines, IA 50319; Illinois Department of Natural Resources, Springfield, IL 62702; Department of Biology, Eastern Michigan University, Ypsilanti, MI 48197; Department of Biology, Bucknell University, Lewisburg, PA 17837

## Abstract

White-nose syndrome (WNS) is a fungal wildlife disease of bats that has caused precipitous declines in certain Nearctic bat species. A key driver of mortality is premature exhaustion of fat reserves, primarily white adipose tissue (WAT), that bats rely on to meet their metabolic needs during winter. However, the pathophysiological and metabolic effects of WNS have remained ill-defined. To elucidate metabolic mechanisms associated with WNS mortality, we infected a WNS susceptible species, the Little Brown Myotis (*Myotis lucifugus*), with *Pseudogymnoascus destructans* (*Pd*) and collected WAT biopsies for histology and targeted lipidomics. These results were compared to the WNS-resistant Big Brown Bat (*Eptesicus fuscus*). A similar distribution in broad lipid class was observed in both species, with ∼60% of total WAT consisting of triacylglycerides (TAGs). We found several baseline differences in WAT chemical composition between species. *M. lucifugus* WAT had significantly higher levels of measured TAGs (∼30%). Higher lipid levels in *E. fuscus* WAT were primarily sphingomyelins and glycerophosphoethanolamines (PEs), along with glycerophospholipids (GPs) dominated by unsaturated or monounsaturated moieties and n-6 (18:2, 20:2, 20:3, 20:4) fatty acids. These differences between *M. lucifugus* and *E. fuscus* may indicate dietary differences that lead to differential “fuel” reserves that are available during torpor. Following *Pd*-infection, we found that perturbation to WAT reserves occurs in *M. lucifugus*, but not in the resistant *E. fuscus*. A total of 36 GPs (primarily PEs) were higher in *Pd*-infected *M. lucifugus*, indicating perturbation to the WAT structural component. In addition to changes in lipid chemistry, smaller adipocyte sizes and increased extracellular matrix deposition was observed in *Pd*-infected *M. lucifugus*. This is the first study to describe WAT lipidomic composition of bats with different susceptibilities to WNS and highlights that recovery from WNS may require repair from adipose remodeling in addition to replenishing depot fat during spring emergence.

## INTRODUCTION

White-nose syndrome (WNS) is a disease of bats caused by the psychrophilic fungus *Pseudogymnoascus destructans* (*Pd*) [1]. It is of particular concern in North America, where it has caused a precipitous decline in certain cave-dwelling species [2]. Since its first detection in 2006 [3], subsequent studies have shown that not all Nearctic bat species are equally susceptible to WNS mortality. In the Midwestern U.S. for example, drastic declines have been observed in *Myotis* species such as the Little Brown Myotis (*Myotis lucifugus*) (>80% in Indiana [4], the Great Lakes Region [5], and Michigan [6]) and Northern long-eared Myotis (*Myotis septentrionalis*) (>90% in Michigan [6], the Great Lakes Region [5], and Arkansas [7]). Declines in other species, including Indiana Myotis (*Myotis sodalis*) [8] and Tricolored Bats (*Perimyotis subflavus*) [9], are also of concern, however, other species such as the Big Brown Bat (*Eptesicus fuscus*), Eastern Red Bat (*Lasiurus borealis*), and Virginia Big-eared Bat (*Corynorhinus townsendii virginianus*) have not experienced the same declines. Within species of concern, intraspecific variation in survival has been observed in populations of *M. lucifugus* [10], possibly influenced by size, as increased fat stores were correlated with increase survival [11]. Understanding the mechanisms and environmental traits driving this variability to WNS-associated mortality among different bat species has been of great interest, as limited fiscal resources are available for management strategies (e.g., habitat protection, hibernacula microclimate manipulation, site treatment) to mitigate the effects of WNS. In addition, elucidating mechanisms of resistance associated with natural bat populations may offer insights into the survivors and potential recovery of species currently experiencing heavy declines [12, 13].

To study the host mechanisms driving disease susceptibility, the effects of *Pd*-infection can be categorized into both cutaneous effects at the site of fungal infection and systemic effects from altered torpor patterns [14, 15] and increased energy expenditure [16]. As bats primarily rely on stored lipid reserves and fatty acid β oxidation to fuel their torpor and rewarming bouts [17], a detailed understanding of fat deposition and altered lipid metabolism is inextricably linked to understanding the WNS process that is needed to frame effective management responses. At early stages of WNS, in addition to changes in blood chemistry [18], perturbations to lipid metabolism are occurring before emaciation is evident [19, 20]. In uninfected *M. lucifugus* and *E. fuscus* from northern populations (Ontario, Canada) during hibernation (mid-January), female bats showed higher hepatic triacylglyceride (TAG) levels compared to males that possibly reflects resource conservation needed for increased spring fecundity [19]. However, after laboratory *Pd*-infection, a non sex-specific increase in some glycerophospholipids (GPs) was observed in *M. lucifugus*, along with a decrease in female TAG levels in both species, while changes in *M. lucifugus* were confined to n-3 TAGs [20]. Although the liver may partially function in lipid storage, hepatic lipid levels are reflective of lipid “shuttling” as it is a primary site for transportation, synthesis, and catabolism [21].

Observation of these early responses to *Pd*-infection motivated further research on the same cohort, which included detailing splenic lipid metabolism as it is a primary site in immune modulation and T and B cell differentiation [20]. Rather than exhibiting immune-specific responses, WNS-susceptible bats (*M. lucifugus*) showed increased oxidative stress that may play a role in downstream deterioration to body condition. Upregulated cutaneous transcription of genes involved in regulating lipid metabolism are also observed in euthermic versus torpid bats and at more advanced stages of WNS [22]. What remains to be determined is 1) how the quality of adipose fat stores may vary among species and regions (e.g., northern vs. southern latitudes) and play a role in survival in addition to total stored fat [11], and 2) what pathological changes may occur within WAT during increased torpor arousals.

In this study, we expand on a previous work that explored physiological variations in two bat species with different susceptibilities to WNS, *M. lucifugus* and *E. fuscus*, after being laboratory-infected with *Pd* and held in identical microclimates [23]. Specifically, as premature exhaustion of stored lipid reserves is a primary contributor to bat mortality, we utilized a targeted lipidomics approach to determine 1) differences in WAT lipid composition between *M. lucifugus* and *E. fuscus* without *Pd*-infection, and 2) changes to the WAT lipid composition in both *M. lucifugus* and *E. fuscus* between a control group and a *Pd*-infection group. We predicted broad species level differences in the WAT chemical composition between *M. lucifugus* and *E. fuscus*. Also, we predicted higher perturbation to lipid reserves in *M. lucifugus* compared to *E. fuscus* after *Pd*-infection. This is the first study to describe WAT composition of bats with different susceptibilities to WNS, provides evidence that *M. lucifugus* and *E. fuscus* may have different “fuel reserves” available to them when entering torpor, and that increased perturbation to adipose metabolism is occurring in *M. lucifugus* compared to *E. fuscus*.

## MATERIALS AND METHODS

### Animal Experiment and Histology

The WAT samples used in this study were collected as part of a captive hibernation experiment performed at Bucknell University from November 2011 to March 2013 [23]. As shown in Table 1 (and Supplementary Table 1), the bats were captured between 5 November 2011 and 17 November 2011 in hibernacula in Michigan or Illinois, USA (*M. lucifugus*) and in Iowa or Illinois, USA (*E. fuscus*). At the time of capture, none of these populations was known to have previous exposure to WNS. The bats were transported while torpid to Bucknell University in Lewisburg, Pennsylvania, USA where they were treated with either phosphate-buffered saline with 0.5% TWEEN-20 (control) or with a suspension of 350,000 Pd conidia (*Pd* exposed). Control and treated bats were separately housed in environmental chambers at 4 C and 95% relative humidity with each species in separate enclosures within the chamber. Following 3, 7, or 13 weeks of hibernation, bats were removed from the chamber and humanely euthanized by isoflurane overdose followed by decapitation. WAT was removed, among other tissues, and snap-frozen in liquid nitrogen. Tissue was stored at −80 C until processed for analysis.

**Table 1.**
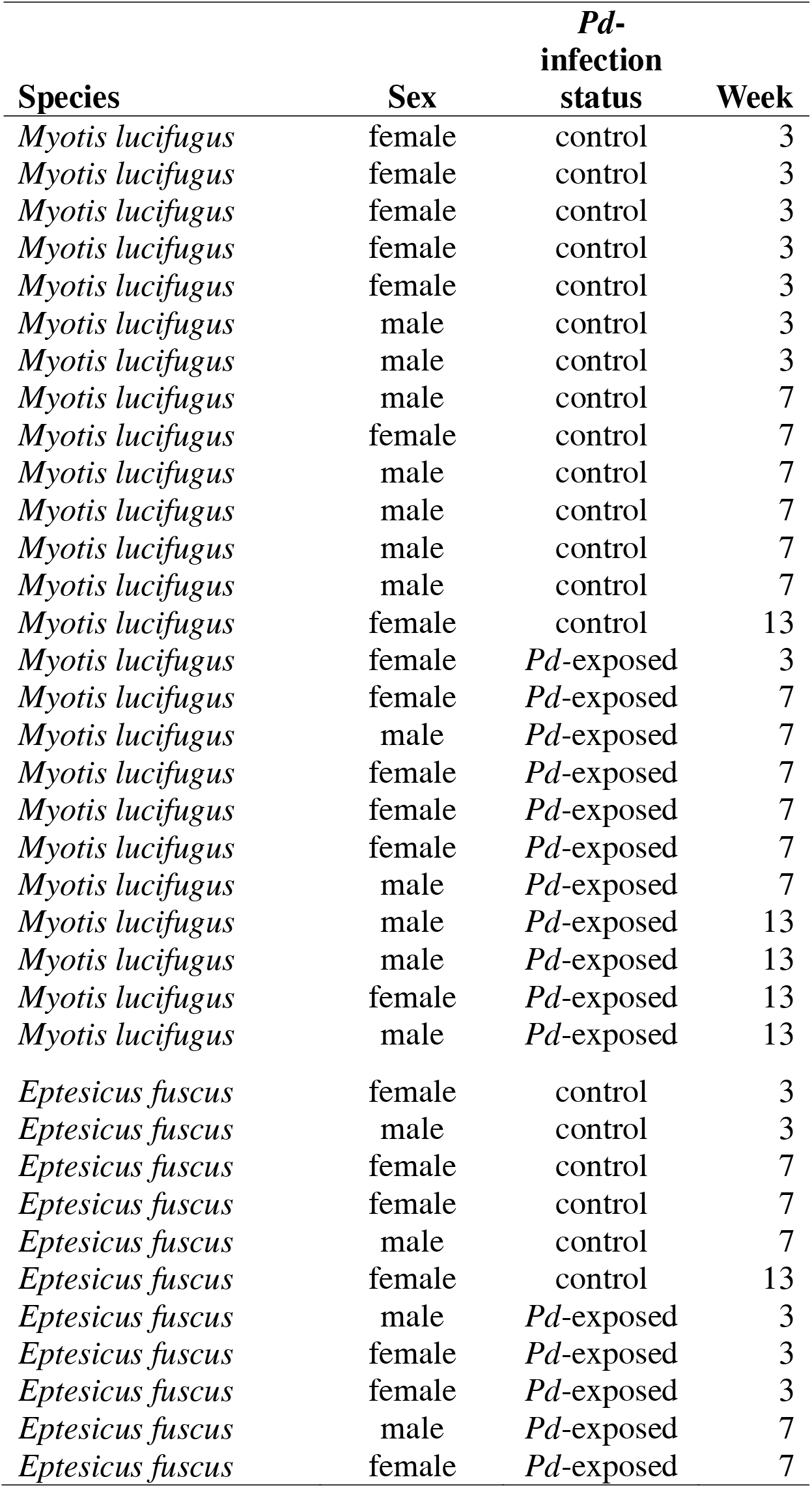
Species, sex, *Pd*-infection status, and week euthanized post-*Pd* exposure for white adipose biopsies used in the current study.

Capture, handling, and sample collection protocols for this study were reviewed and approved by the Bucknell University IACUC (protocol #DMR-12), and the US Fish and Wildlife Service. In the states of Illinois and Iowa, research collections were conducted by state wildlife officials and on non-endangered bats; thus, numbered permits were not required or issued. In Michigan, research was conducted under Scientific Collector’s Permits #SC1448 to DMR.

### Microscopy and Histology

Flash-frozen WAT samples were immediately stored at −80 °C until fixation. Samples were kept on ice for 20 minutes and then placed in a tube with a 1:10 volume of fixative-1 solution (20% formaldehyde, 2% glutaraldehyde in phosphate buffer saline pH 7.4) at room temperature for 30 minutes. Finally, samples were transferred to a 10% formalin solution and incubated for 2-3 hours at room temperature for final fixation, then transferred into 70% ethanol followed by paraffin embedding. Formalin-fixed paraffin-embedded (FFPE) sections were used for hematoxylin and eosin (H&E) staining using the standard protocol. The BX-63 microscope mounted with DP 28 digital camera (Olympus, MA, USA) was used to perform light microscopy on H&E-stained adult male bat WAT sections (*M. lucifugus* control n = 6 and Pd-infected n = 5). Digital images were acquired using cellSens Entry v1.15 (Olympus, MA, USA) software at a fixed setting for both control and infected groups. Histological assessment of adipocyte size was estimated using the Adiposoft v1.16 plug-in for ImageJ [24]. The automated settings (min size = 5, max size = 60) were used, and then the output was checked manually. WE compared the average adipocyte cell sizes using an unpaired t-test in GraphPad Prism 9.2.0 (GraphPad Software, La Jolla, CA).

### Lipid Extraction and Profiling

Fisher Optima^TM^ grade reagents were used for sample preparation and analysis (Fisher Scientific, Hanover Park, IL). EquiSPLASH® LIPIDOMIX (Avanti Polar Lipids Inc., Alabaster, AL, USA) was used for the internal standard and NIST plasma Standard Reference Material (SRM) 1950 was prepared as a quality control.

To assess the linearity of different lipids for different tissue amounts, we collected a 62.9-mg biopsy of WAT from an individual female adult parous EPFU and extracted the lipid content into methanol (300 μl) with internal standards (5 μl, EquiSPLASH® LIPIDOMIX) according to manufacturer’s instructions. Samples were left at −80 °C overnight, and on the following day, they were placed on dry ice and homogenized. After homogenization, samples were vortexed, incubated on ice (10 min), and cold chloroform (600 μl) was added. Samples were then vortexed, incubated on ice (10 min), and water (300 μl) was added. Samples were then vortexed, incubated on ice (10 min), and centrifuged for 5 min (1000 rpm, 4 °C). The bottom layer was removed and dried via lyophilization. The sample was reconstituted in isopropanol:acetonitrile (IPA:ACN 75:25, 200 μl) and diluted 10-fold for a 5-point curve.

The optimal tissue weight for analysis fell between ∼0.5 – 10 mg, at which we see a loss of linearity (primarily in TAGs) with tissue weights over 10 mg (Supplementary Fig 1). Excellent linearity (R^2^ > 0.99, ePE R^2^ = 0.97) was observed between 0.62 – 6.29 mg tissue weights; however, for the broad classes that are found in naturally lower proportion in WAT (e.g., CERs, CEs, acylcarnitines, and minor GPs and LGPs), several compounds will not be detected in smaller tissue biopsies, which needs to be considered if attempting to analyze small micro-adipose samples from nonlethal sampling [25]. Although, it is important to note that these results are highly instrument specific and newer platforms offer improved sensitivity and dynamic range for detecting lower concentration compounds in biological samples.

Samples were injected (2 μl) and analyzed using a Waters Acquity Ultra Performance Liquid Chromatography (UPLC) (CSH C18 1.7 μm, 2.1 x 100 mm column) and a Xevo^®^ G2 quadrupole time-of-flight (QTOF) mass spectrometry (MS) (Waters, Milford, MA, USA). Positive and negative electrospray ionization (ESI) data-independent modes were used for data acquisition. Leucine enkephalin ([M+H]^+^ = 556.2771, [M-H]^-^ = 554.2615) was used as Lock-Spray^®^. Operating conditions for ESI were: capillary voltage 3.0 kV, cone voltage 30 V, desolvation temperature 500°C, and desolvation gas flow 1000 L/Hr. Mobile phases consisted of the following: solvent A (water/0.1% formic acid [FA]/10 mM ammonium formate), solvent B (ACN/0.1% FA), and solvent C (IPA/0.1% FA). The gradient was 8.0 min (solvent A 30%, B 34%, C 36%), 0.5 min 20% (solvent A 0%, B 10%, C 90%), and 2.5 min (solvent A 30%, B 34%, C 36%) at a flow rate of 0.45 ml/min and the column temperature maintained at 65 °C.

Our targeted lipid profiling assay has been previously described with lipid classes and multiple reaction monitoring (MRM) transitions listed [20, 26]. Briefly, ∼ 10 mg of tissue was homogenized as above, with tissue disruption after a 12-hr period at −80 °C, lipid isolation/purification using a liquid-liquid extraction, and lyophilization (Figure 1). The dried extract was reconstituted in 200 μL of extraction buffer (isopropanol) and filtered using a 0.2 μM microcentrifuge filter. The supernatant was transferred to MS vial for LC-MS analysis. As WAT is inherently low in both water and protein content, we normalized to weight rather than protein concentration to minimize the amount of sample needed for normalization and the contribution to the weight from water should be minimal. Quality control samples of both a pooled sample and the NIST SRM 1950 plasma mix were run every 10 samples. The SIL-30 AC auto sampler was kept at 15°C (Shimazdu, Columbia, MD, USA). A 5 µL portion was injected and analyzed (Xbridge amide column, 3.5 µm, 4.6 × 100 mm) (Waters, Milford, MA, USA) using a Sciex QTRAP 5500 Mass Spectrometer (Sciex, Framingham, MA, USA). Lipids were detected by multiple reaction monitoring (MRM) transitions in both positive and negative ionization mode. Operating conditions were: temperature□=□550□°C, nebulizing gas□=□50 and heater gas□=□60, curtain gas□=□30, CAD gas□=□medium, ion spray voltage□=□5.5□kV in positive mode and −4.5□kV in negative mode. Mobile phases consisted of the following: solvent D (95% acetonitrile/5% water with 10 mM ammonium acetate) and solvent E (50% acetonitrile/50% water with 10 mM ammonium acetate). The mobile phase was initially 100% solvent D, then a gradient of 3.0 min (solvent D 99.9%, E 0.01%), 3.0 min (solvent D 94%, E 6%), 4.0 min (solvent D 25%, E 75%), 6.0 min (solvent D 0%, E 100%), and then 6.0 min equilibration back to 100% solvent D at a flow rate of 0.7 ml/min and the column temperature maintained at 35 °C.

**Figure 1.**
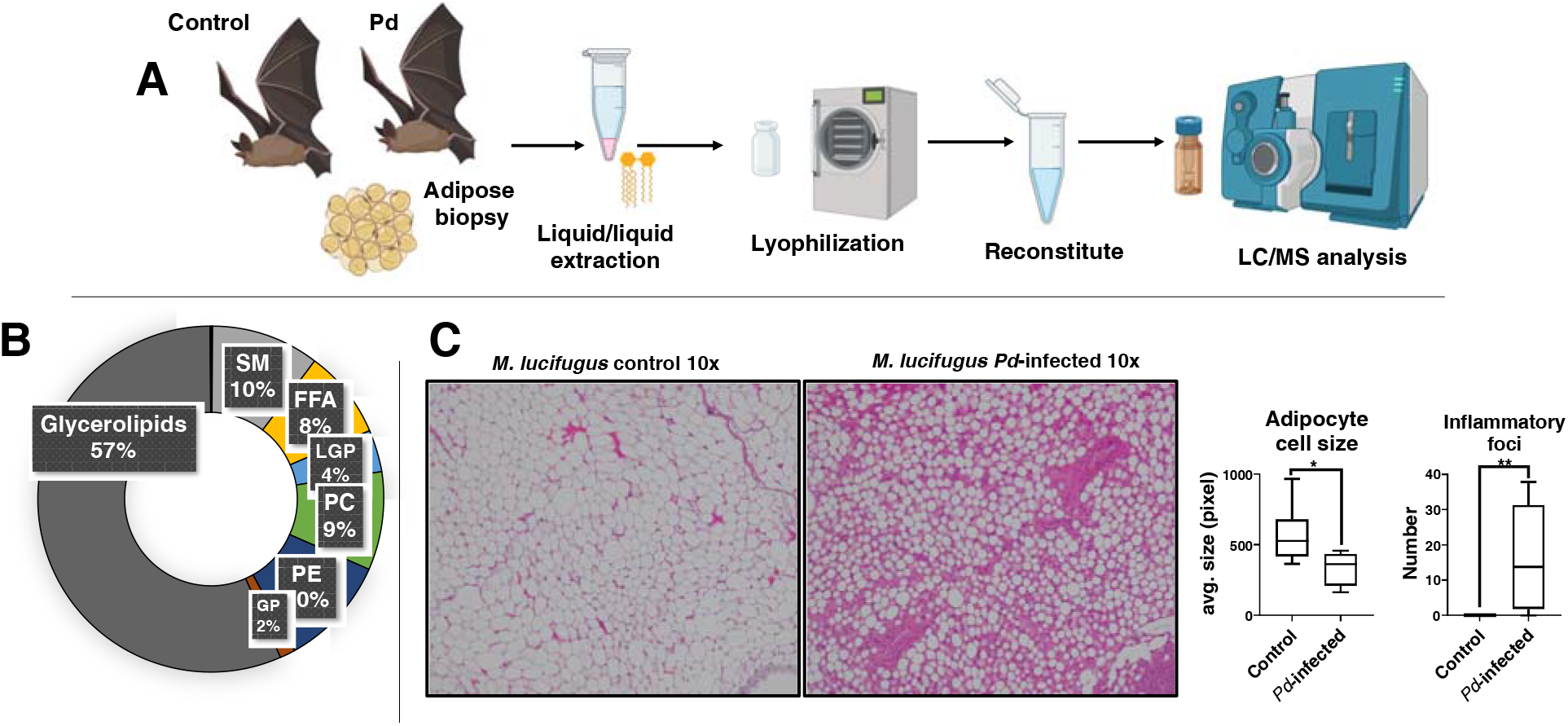
**A)** General schematic for tissue sampling and analytical workflow. **B)** Donut plot depicting the chemical composition of *Myotis lucifugus* (n=14) white adipose tissue (WAT) in the control group. The overall proportion is similar to what is observed in other mammals. **C)** H&E staining of WAT sections (n=5) from *M. lucifugus* showing decreased adipocyte size in the *Pd*-infected group. Bats used in the current study also had a higher number of inflammatory foci in the *Pd*-infected group. (*P ≤ 0.05; lines from top to bottom represent max value, 75th percentile, median, 25th percentile, min value.)

### Data Analysis

Untargeted raw data files were imported into MassLynx v.4.1 (Waters Corporation, Milford, MA) and aligned used Progenesis QI (Nonlinear Dynamics, Newcastle, UK). Normalization was performed using a software chosen QC chromatogram as an alignment reference and normalizing to the “normalize to all compounds function”. For lipids of interest that were identified from the below targeted dataset, we visually inspected the *m/z* and MSe fragmentation patterns to confirm identities, and then standard curves were plotted in GraphPad Prism 9.2.0 the obtain R^2^ values (GraphPad Software, La Jolla, CA). For the targeted lipidomics dataset, the qlm was imported into MultiQuant v 2.0 (Sciex, Framingham, MA, USA), peak areas were visually inspected, and the dataset was exporting to excel. We first removed lipids in the pooled QC sample with a coefficient of variation >30%. We compared the control groups for *M. lucifugus* (n=14) and *E. fuscus* (n=6) (false-discovery rate [FDR] corrected P < 0.05 using a standard Benjamini-Hochberg procedure based on R p.adjust(raw.pvals.”fdr”)) for species level differences. To determine if sex differences between uninfected *M. lucifugus* adult females vs. males (n=5 for both groups) could influence the results of infection, we screened for statistically significant lipids (FDR corrected P < 0.05). The sample size in the current study did not permit sex comparisons within *E. fuscus.* To determine disease effects for *M. lucifugus* (n=14 control; n=11 *Pd*-infected) and for *E. fuscus* (n=6 control; n=5 *Pd*-infected), we used the FDR corrected P ≤ 0.15 as a cutoff which corresponded to lipids with the highest fold changes. The sample size in the current study did not permit sex comparisons within *E. fuscus.* Lipid profiles were screened in SAS 9.4 (SAS, Cary, NC) and by using the t test feature in MetaboAnalyst 5.0 [27, 28]. MetaboAnalyst 5.0 was also used to construct heatmaps and box-and-whisker plots were generated in GraphPad Prism 9.2.0 (GraphPad Software, La Jolla, CA).

## RESULTS

We measured the abundance of various lipid groups in the WAT tissues of *M. lucifugus* and *E. fuscus*, with and without *Pd* exposure, predicting that we would find differences in the WAT chemical composition that depended on species, WNS-susceptibility, and sex. Several lipid groups were measured, which we broadly grouped based on function and similarity, including the glycerolipids (predominantly TAGs, but also a minor diacylglyceride [DAG] and monoacylglyceride [MAG] component), sterols (predominantly cholesterol with a minor cholesteryl ester [CE] component), the ceramides (CERs) (including minor amounts of hexosylceramides [HexCers], lactosylceramides [LCERs], and dihydroceramides [DCERs]) with sphingomyelins (SMs) separated out, acylcarnitines, free fatty acids (FFAs), the phosphatidylcholines (PCs) (including ether-linked phosphatidylcholines [ePCs]), the phosphatidylethanolamines (PEs) (including ether-linked phosphatidylethanolamines [ePEs]), and the rest of the minor GPs (phosphatidylglycerols [PGs], phosphatidic acids [PA], phosphatidylinositols [PI], and phosphatidylserine [PS]) or lysoglycerophospholipid (LGP) groups combined (lysophosphatidic acids [LPA], lysophosphatidylcholines [LPC], lysophosphatidylinositols [LPI], and lysophosphatidylethanolamines [LPE]).

Overall, a similar distribution was observed to that described in humans [29], with the WAT composition in both *M. lucifugus* and *E. fuscus* is dominated by TAGs and a smaller proportion of glycerolipids (57% and 52% combined glycerolipid respectively), followed by PCs, PEs, FFAs, SMs, and other GPs (Figure 1). Sterols made up 0.1% of the WAT proportion (95% of which was cholesterol), and the ceramides and acylcarnitines were collectively <0.1%. This small proportion of ceramides differs from human WAT, which has a 4.3% ceramide fraction [29]. An opposite trend in the glycerolipid proportion of WAT was observed in *Pd*-infected *M. lucifugus* and *E. fuscus*, as *M. lucifugus* had a minor decrease in glycerolipid proportion (49%) (Supplementary Fig 2). An increase in the WAT glycerolipid proportion was observed in the *E. fuscus* cohorts (control = 52% and increased to *Pd*-infected = 69%), possibly due to the increased average torpor bout length observed in the *Pd*-infected group compared to the control group (Supplementary Fig 2) [23]. WAT remodeling was observed in *Pd*-infected *M. lucifugus*, in which a smaller adipocyte cell size with a large effect size (P = 0.05; Cohen’s *d* = 1.38) was observed in tissue sections along with increased ECM (Figure 1).

The targeted lipid dataset was further analyzed by broad group, including glycerolipids, GPs and SMs, FFAs, and low abundance compounds (i.e., HexCers, LCERs, DCERs, PAs, LPAs, CEs, and acylcarnitines). We first looked for sex differences between adult male and female *M. lucifugus,* the species for which we had a sufficient sample size to examine potential differences. Upon finding no significant differences in the WAT chemical composition between adult male and female *M. lucifugus*, we lumped the sexes for further analysis. Importantly, although total proportion among bat species are similar, the chemical composition between *M. lucifugus* and *E. fuscus* WAT differs dramatically. We found 155 statistically significant differences in lipids between species (Figure 2, Supplementary Table 1). Higher lipid levels in *E. fuscus* WAT were primarily in the broad classes SMs and PEs. Only five glycerolipids (DAG [18:1/18:1], [18:1/18:2], [18:1/20:1] and TAG 54:5-FA18:2, 54:6-FA18:2) were higher in *E. fuscus* along with nine PCs. Higher GP levels in *E. fuscus* WAT were dominated by compounds containing unsaturated or monounsaturated moieties along with potential n-6 (18:2, 20:2, 20:3, 20:4) fatty acids that comprised ∼80% of the GP fraction. Most of lipids found at a statistically significant level were higher in *M. lucifugus* WAT, including 83 TAGs. Increased TAG levels in control *M. lucifugus* spanned the range of TAGs included in our assay, both in terms of total carbon number (48 – 56 carbons) and saturation (unsaturated – 8 double bonds). Compared to *E. fuscus*, lipids that are in higher concentration in *M. lucifugus* (other than TAGs) tended to contain potential n-3 (18:3, 20:5, 22:5, 22:6) fatty acids, showing an even 1:1 ratio of n-6/n-3 ratio. As mammals cannot synthesize these lipids, differences between these two species were likely due to diet. Overall, these results indicated *M. lucifugus* may enter torpor with a more highly concentrated WAT compared to *E. fuscus*.

**Figure 2.**
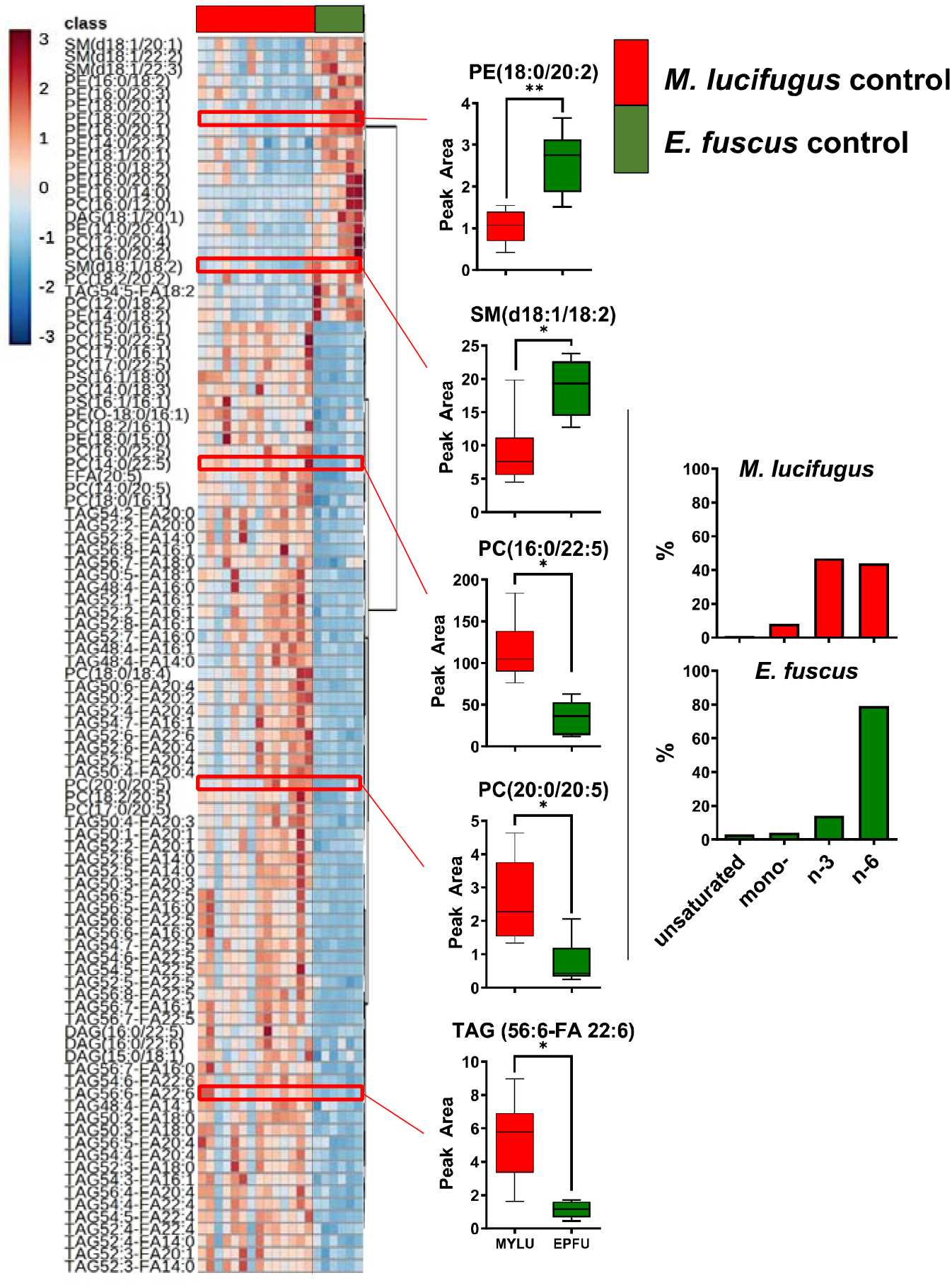
Species level differences in lipid compounds in white adipose tissue (WAT) between the *Myotis lucifugus* (n=14) and *Eptesicus fuscus* (n=6) control groups. Although broad lipid class proportions are similar between the species, there are distinct differences in the WAT chemical composition, which may have important downstream implications for over winter energy reserves. Box and whisker plots of some select lipids show a trend for increased concentration of lipids with n-6 constituents in *E. fuscus* converse to more highly unsaturated n-3 compounds in *M. lucifugus*. (*FDR corrected P ≤ 0.01; ** FDR corrected P ≤ 0.001, lines from top to bottom represent max value, 75th percentile, median, 25th percentile, min value., P value and fold change are listed in Table S2)

Comparing the uninfected and *Pd*-infected groups within each species, we did not find any significant differences in the glycerolipids or in low abundance compounds (Supplementary Table 3). The remaining lipids were ranked by P value, with the top 38 lipids having an FDR corrected P value = 0.15 (Table 2). A total of 36 lipids were higher in *Pd*-infected *M. lucifugus* compared to uninfected *M. lucifugus* including 8 PCs, 1 PG, and 2 PSs (PS [18:0/18:1], [18:0/18:2]), with the remainder as PEs (Figure 3, Supplementary Fig 3). Functionally, these compounds are structural in nature as they comprise cellular membranes, with PEs also being found in high levels in mitochondria [30]. These observations on WAT TAG concentration, together with the increasing concentration of GPs, correlates with the histological observation that WAT remodeling is occurring along with increased ECM deposition in *M. lucifugus* during *Pd*-infection but not in *E. fuscus*.

**Figure 3.**
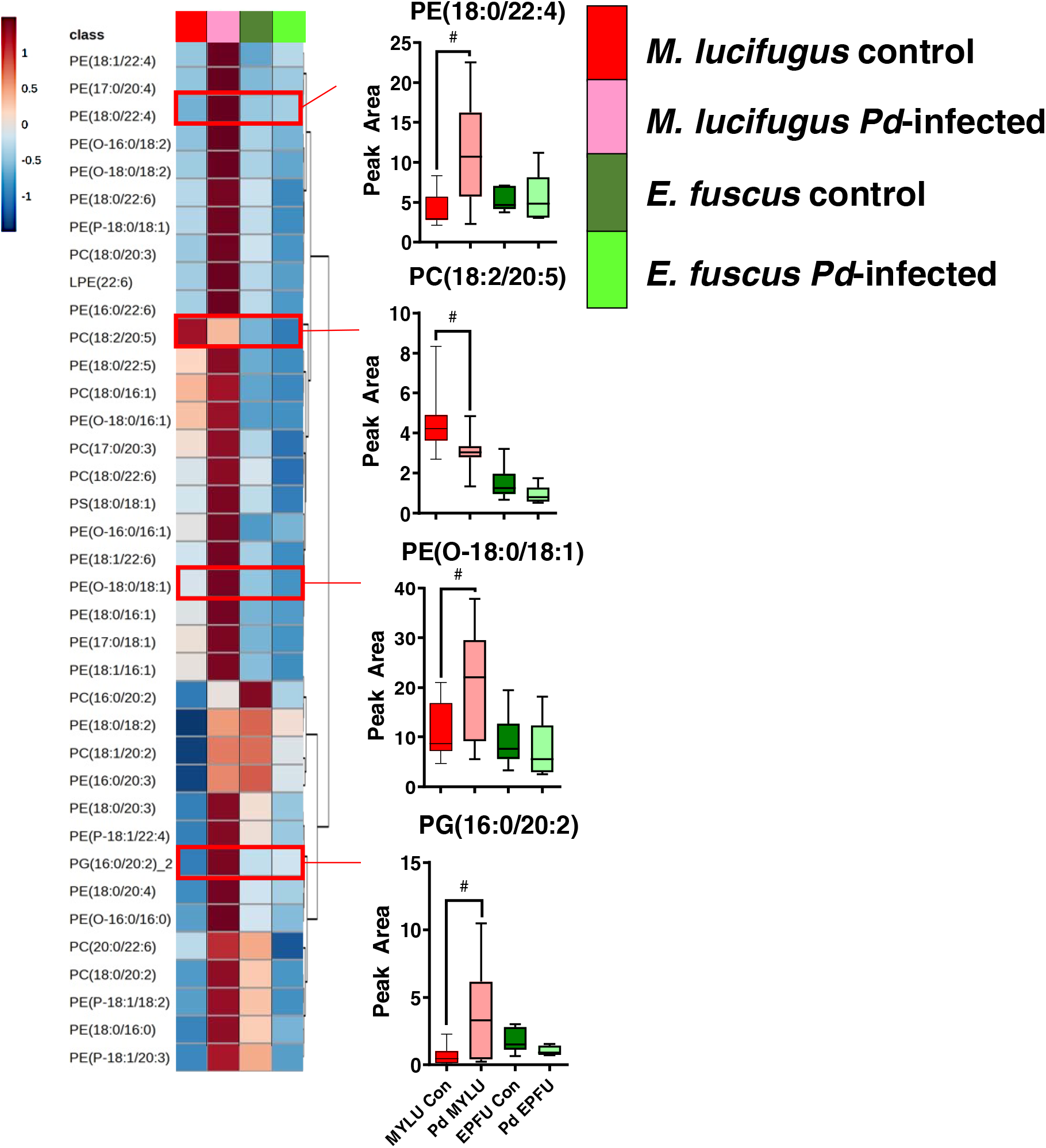
Differences in lipid compounds in white adipose tissue (WAT) between the *Myotis lucifugus* control group (n=14) and *Pd*-infected group (n=11). Values for *Eptesicus fuscus* groups (n=6 control, n=5 *Pd*-infected) are provided as a comparison. Disease level changes were only observed in glycerophospholipid levels, primarily phosphatidylethanolamines, indicating a greater contribution from structural membranes or increased mitochondrial numbers. (#FDR corrected P = 0.15, lines from top to bottom represent max value, 75th percentile, median, 25th percentile, min value., an expanded heatmap with individuals used in study supplied in Fig S3, P value and fold change are listed in Table 2)

**Table 2.** Top Ranked Lipids Found in White Adipose Tissue of *Myotis lucifugus* Comparing Control Group vs. *Pd*-infection Group (normalized peak area Mean ± Std Dev).

|  | *M. lucifugus*<br>Control | *M. lucifugus*<br>*Pd*-infected | Fold<br>Change | P value | FDR |
| --- | --- | --- | --- | --- | --- |
| LPE(22:6) | 0.42±0.40 | 1.02±0.72 | 2.4 | 0.0166 | 0.154 |
| PC(16:0/20:2) | 1.54±0.67 | 2.48±0.98 | 1.6 | 0.0108 | 0.152 |
| PC(17:0/20:3) | 1.27±0.67 | 1.98±0.70 | 1.6 | 0.0189 | 0.154 |
| PC(18:0/16:1) | 52.71±14.54 | 76.91±31.06 | 1.5 | 0.0188 | 0.154 |
| PC(18:0/20:2) | 0.84±0.45 | 1.99±1.17 | 2.4 | 0.0033 | 0.152 |
| PC(18:0/20:3) | 17.44±7.44 | 35.32±24.96 | 2.0 | 0.0203 | 0.154 |
| PC(18:0/22:6) | 10.65±7.79 | 20.02±8.51 | 1.9 | 0.0100 | 0.152 |
| PC(18:1/20:2) | 1.73±0.92 | 3.28±1.67 | 1.9 | 0.0084 | 0.152 |
| PC(18:2/20:5) | 4.48±1.42 | 3.08±0.91 | 0.7 | 0.0103 | 0.152 |
| PC(20:0/22:6) | 0.53±0.31 | 0.92±0.43 | 1.7 | 0.0161 | 0.154 |
| PE(16:0/20:3) | 2.14±1.49 | 4.64±2.79 | 2.2 | 0.0097 | 0.152 |
| PE(16:0/22:6) | 5.01±3.71 | 12.61±8.08 | 2.5 | 0.0054 | 0.152 |
| PE(17:0/18:1) | 5.26±3.18 | 10.46±4.57 | 2.0 | 0.0033 | 0.152 |
| PE(17:0/20:4) | 7.28±5.09 | 14.48±8.73 | 2.0 | 0.0188 | 0.154 |
| PE(18:0/16:0) | 1.19±0.63 | 2.37±1.40 | 2.0 | 0.0109 | 0.152 |
| PE(18:0/16:1) | 4.11±2.58 | 9.58±6.04 | 2.3 | 0.0065 | 0.152 |
| PE(18:0/18:2) | 270.06±128.64 | 498.69±254.68 | 1.8 | 0.0088 | 0.152 |
| PE(18:0/20:3) | 12.84±8.96 | 31.73±18.93 | 2.5 | 0.0037 | 0.152 |
| PE(18:0/20:4) | 312.84±222.04 | 600.15±315.44 | 1.9 | 0.0155 | 0.154 |
| PE(18:0/22:4) | 4.75±2.90 | 11.29±6.78 | 2.4 | 0.0041 | 0.152 |
| PE(18:0/22:5) | 50.61±23.37 | 89.34±50.61 | 1.8 | 0.0204 | 0.154 |
| PE(18:0/22:6) | 15.75±11.67 | 40.40±33.88 | 2.6 | 0.0205 | 0.154 |
| PE(18:1/16:1) | 53.92±34.86 | 110.23±60.78 | 2.0 | 0.0090 | 0.152 |
| PE(18:1/22:4) | 11.19±8.43 | 21.80±11.44 | 1.9 | 0.0156 | 0.154 |
| PE(18:1/22:6) | 28.42±24.10 | 65.09±43.17 | 2.3 | 0.0148 | 0.154 |
| PE(O-16:0/16:0) | 0.51±0.23 | 0.93±0.48 | 1.8 | 0.0095 | 0.152 |
| PE(O-16:0/16:1) | 1.40±0.73 | 2.39±1.17 | 1.7 | 0.0187 | 0.154 |
| PE(O-16:0/18:2) | 2.05±1.37 | 4.23±2.68 | 2.1 | 0.0165 | 0.154 |
| PE(O-18:0/16:1) | 8.92±2.37 | 13.65±5.61 | 1.5 | 0.0103 | 0.152 |
| PE(O-18:0/18:1) | 11.32±5.29 | 20.31±10.01 | 1.8 | 0.0095 | 0.152 |
| PE(O-18:0/18:2) | 30.47±16.63 | 58.32±30.13 | 1.9 | 0.0085 | 0.152 |
| PE(P-18:0/18:1) | 31.55±21.29 | 70.27±50.76 | 2.2 | 0.0187 | 0.154 |
| PE(P-18:1/18:2) | 36.65±26.04 | 89.63±66.03 | 2.4 | 0.0131 | 0.154 |
| PE(P-18:1/20:3) | 11.80±10.04 | 26.88±16.35 | 2.3 | 0.0107 | 0.152 |
| PE(P-18:1/22:4) | 17.41±15.66 | 46.74±35.92 | 2.7 | 0.0131 | 0.154 |
| PG(16:0/20:2)_2 | 2.99±3.04 | 11.03±10.05 | 3.7 | 0.0106 | 0.152 |
| PS(18:0/18:1) | 12.45±10.07 | 26.26±16.50 | 2.1 | 0.0188 | 0.154 |
| PS(18:0/18:2)_2 | 0.53±0.18 | 0.85±0.41 | 1.6 | 0.0194 | 0.154 |

## DISCUSSION

During *Pd*-infection, bats in susceptible species undergo increased torpor arousals during winter months and prematurely exhaust fat stores and die due to emaciation [14, 15]. As not all bats are equally susceptible to WNS associated mortality, a key research priority has been deciphering the traits driving this differential susceptibility among species. While total fat stores have been the focus of previous studies [31], another factor that has been less explored is potential differences in prehibernation WAT quality and pathological changes during WNS. Here, we found that while ∼60% of total WAT is comprised of TAGs and other glycerolipids associated with energy storage, a large proportion of WAT is comprised of other lipid classes, such as GPs and SMs. In *M. lucifugus*, ∼30% of measured TAGs were significantly higher compared to *E. fuscus*, which may indicate increased energy density. For GPs, significantly higher compounds found in *E. fuscus* WAT had n-6 fatty acid moieties, whereas n-3 dominated in *M. lucifugus*. Compared to hibernating bats without infection, *M. lucifugus* infected with *Pd* showed WAT remodeling including increased presence of ECM, decreased adipocyte size, and increased GP levels.

The most striking difference observed in this study was the species level differences observed in the WAT composition between *M. lucifugus* and *E. fuscus*. This is not surprising as these are different species and they consume different types of insects [32, 33]. The ratio of depot fat to lean mass of hibernators peaks in the fall before they reduce their metabolic rate and begin torpor bouts, so the “snapshot” of WAT composition presented in this work represents the metabolic fuel that would be available for periodic rewarming during winter and sustain them for spring foraging and female reproduction [34]. Interestingly, we found several glycerolipids (82 TAGs, representing 32% of all TAGs measured, and 6 DAGs) in higher concentration in *M. lucifugus* compared to *E. fuscus*, while only 2 TAGs and 3 DAGs were found higher in *E. fuscus*. In addition to species-level differences in adipocyte biology being well recognized [35], fat composition and lipid content have also been linked to hibernation strategies between different bat species [36]. Having a significantly higher TAG density may indicate *M. lucifugus* has more energy dense WAT compared to *E. fuscus*; however, it is also possible the difference in torpor patterns between these species may influence the observed WAT TAG density [23]. Another interesting aspect is the significantly higher amounts of n-3 GPs found in *M. lucifugus*, while *E. fuscus* had predominantly higher levels of n-6 GPs.

Several studies have highlighted that not all fats are equal, in that during hibernation there is a selective retention of lipids based on saturation and that the n-6/n-3 ratio affects torpor quality (see [37] for review). Even in animals with dramatically different hibernation patterns (captive garden dormice [*Eliomys quercinus*] vs. free-ranging brown bears [*Ursus arctos*]), elevated n-3 plasma GPs were observed suggesting a common hibernation lipid phenotype [38]. A potential mechanism for these observations is that increased membrane fluidity may be needed to maintain function, such as sarcoendoplasmic reticulum calcium ATPase activity [39]. Further research may elucidate if perturbation to n-6/n-3 membrane ratios may play a role in differential torpor patterns between *M. lucifugus* and *E. fuscus* during WNS.

The basic component of WAT is the adipocyte, which is a unilocular structure consisting of stored TAGs (and other glycerolipids and cholesteryl esters) surrounded by a thin rim containing the cell membrane, cytoplasm, and organelles. The adipocytes are surrounded by a structural ECM that together forms a type of connective tissue. Within WAT, the adipocytes are metabolically active, whereas the ECM is composed of noncellular components, including collagens, laminins, fibronectin, and proteoglycans and not generally considered metabolically active [40]. During periods of torpor and low prey availability, adipocytes play a vital role as they release FFAs from stored TAGs providing fuel through fatty acid β oxidation [41], and during the hibernation period and periodic arousals ∼85% of total WAT is consumed [42]. The chemical composition of WAT can change during these periods of differential energy output (e.g., exercise [43] or winter torpor) and also other factors, such as region of the body [29, 44] or diet [45]. In addition, WAT remodeling can be associated with an aging or pathogenic state. Patients with cancer-induced cachexia show decreased adipocyte size along with increased cell atrophy, fibrosis, and ECM deposition [46]. Pre-hibernation WAT remodeling has also been described in a food-storing hibernator (Syrian hamster, *Mesocricetus auratus*, [47]). In torpor and early arousal, the thirteen-lined ground squirrel (*Ictidomys tridecemlineatus*) shows suppressed inflammatory cytokine and matrix metalloproteinase activity in WAT suggesting that tissue remodeling is not typical during this period in a nonpathogenic state [48]. A hallmark of WNS is the premature exhaustion of fat stores, so a reduction in adipocyte size or number comprising WAT would be expected. In this regard, our observation of a decrease in adipocyte cell size and an increase in ECM indicates both histological and subsequent metabolic remodeling of WAT, reflected through changes in lipid compounds. Because a dynamic balance between adipocytes and the composition of the ECM within WAT is essential for its normal metabolic function and torpor [40], further studies are required to characterize the ECM changes and their implications in adipocyte metabolism between species during WNS.

Inherent to understanding the WNS process is determining metabolic changes that are occurring in resistant vs. susceptible bat species during *Pd*-infection [49, 50]. These changes will occur in tandem with the metabolic changes associated with normal torpor arousal, which has presented some difficulty in teasing out the effects from infection and interbout arousals. One approach used an RNA-Seq method to analyze transcript levels in wing tissue with/without *Pd*-infection during periods of torpor or euthermy in *M. lucifugus* [51, 52]. Interestingly, two of the most consistently upregulated genes due to local *Pd*-infection were involved in lipid metabolism, including *FFAR2* (free fatty acid receptor 2) and *PTGS2* (cyclooxygenase). *FFAR2* was highly upregulated due to both euthermy and *Pd*-infection. It is activated by short chain fatty acids (SCFAs) acetic acid, propionic acid (3 carbon fatty acid), and butyric acid (4 carbon fatty acid) that are primarily microbially produced and play a major role in adipocyte metabolism [53]. In addition to being directly metabolized for energy, SCFAs can influence adipocyte lipolysis, adipogenesis, and browning of adipose tissue. SCFAs can may also modulate inflammation by attenuating cytokine levels; however, much of these data are derived from *in vitro* experiments and their *in vivo* response in wild bat populations remains to be determined. A more direct correlation to inflammation was identified as increased transcription of *PTGS2*, which encodes for cyclooxygenase-2 (COX-2: EC 1.14.99.1) and is involved in the oxygenation of arachidonic acid (FFA 20:4) to the eicosanoid, prostaglandin H_2_ (PGH_2_). These downstream metabolites of arachidonic acid are primary drivers in inflammation, which has made them interesting candidates to study in WNS. PGH_2_ can also be converted to PGI_2_ (prostacyclin), an unstable lipid that is measured in the body as 6-keto-PGF1α. Levels of 6-keto-PGF1α were significantly increased in the spleen tissue of *M. lucifugus* at early stages of WNS, however, it plays a more important role in vasodilation and platelet formation rather than inflammation [20]. Analysis of the metabolites involved in the eicosanoid pathway at more advanced stages of WNS and the presence/severity of inflammation between species and torpor vs. arousals remains an area where additional data are needed.

Finally, from a more technical standpoint, there has been interest in developing non-lethal sampling techniques to elucidate the WNS process to minimize both the number of animals removed from the wild and possible confounding effects from laboratory infection [54]. Non-lethal sampling methods from wild bat species have also been proposed for other types of studies, including viral monitoring and discovery [55]. Sample collection has typically included wing biopsies/wipes [56, 57], blood [58], saliva [59], and fecal samples [60]. In addition, adipose tissue biopsies have been used to measure changes in lipid content by fatty acid profiling via chemical derivatization to fatty acid methyl ester (FAME) analysis [25, 45]. Although derivatization can increase sensitivity that is useful for low tissue masses, a limitation inherent to FAME analysis is that structural information is lost after fatty acid cleavage from their headgroup (e.g., glycerol, sphingosine, cholesterol), and fractionation of lipid subclasses prior to analysis is a tedious process [61]. While this loss of structural information is not as consequential when analyzing homogenous chemical mixtures that have similar function, such as TAGs that will be used for fatty acid β oxidation or hydrocarbons for energy combustion, complications in biological interpretation may arise when dealing with heterogeneous mixtures with varied biological function. We found (in addition to others [29]) that the WAT has a significant fraction of GPs and sphingolipids involved in cellular structure and ECM maintenance [62] that would not been detected using fatty acid profiling. Here, we found that total lipid profiling (i.e., lipidomics) was capable of analyzing small tissue masses that could be collected in a nonlethal manner and offer a level of sensitivity to detect disease severity in wild populations. Although lipid structural integrity is maintained during this analysis, the complexity of data interpretation does increase as a higher number of analytes are detected. During a typical lipidomics experiment, ∼1,000 – 10,000 compounds are detected (depending on targeted vs. untargeted analysis) vs. ∼5 – 20 features measured during fatty acid profiling (assuming mammalian samples).

## CONCLUSIONS

Hibernating bats are unique as the demands for both flight and enduring long periods of extreme temperatures with low prey availability have shaped their natural metabolic and immune responses. The introduction of WNS has added an additional metabolic strain in Nearctic species leading to increased energy expenditure beyond their typical capacity. However, difficulty in studying wild bat populations has hampered deciphering these responses compared to other mammals that are easily bred and housed in laboratory settings. Hence, even well after a decade of the discovery of WNS in North America and identification of the causal agent, the pathophysiological effects during the disease process and how they differ between species are still unclear. We highlighted differences in WAT lipid composition between *M. lucifugus* and *E. fuscus* that may have implications on pre-hibernation strategies between these species. As expected, histological and chemical changes are only detected in the WNS sensitive *M. lucifugus* following *Pd*-infection. Perturbation to the WAT in *Pd*-infected *M. lucifugus* manifested as smaller adipocyte size, increased ECM deposition, and increased in GP levels. We did not observe significantly decreased levels of TAGs in *Pd*-infected *M. lucifugus* that would likely be associated with free-ranging populations, which is in line with the lack of change in body condition [23], and may be an artifact of study duration. As previous studies have focused solely on total fat loss during WNS, these studies have shown that additional metabolic stress is occurring in bats at early stages of disease [19, 20] and that the effects from *Pd*-infection to adipose stores includes ECM deposition that may impact recovery periods beyond simply replenishing fat stores during spring emergence.

---

**Supplementary Table 1.** Metadata of bats used in the current study.

**Supplementary Table 2.** Lipids that are significantly different *Myotis lucifugus* (n=14) and *Eptesicus fuscus* (n=5) that were used as the control groups.

(Mean ± Std. Dev., P Value and FDR P Value calculated in MetaboAnalyst 5.0) MRM transitions have been previously published and are available here: [20]

**Supplementary Table 2.** Differences in lipid compounds in white adipose tissue (WAT) between the *Myotis lucifugus* control group (n=14) and *Pd*-infected group (n=11).

Values for *Eptesicus fuscus* groups (n=6 control, n=5 *Pd*-infected) are provided as a comparison.

(Mean ± Std. Dev., P Value and FDR P Value calculated in MetaboAnalyst 5.0)

**Supplementary Figure 1.**
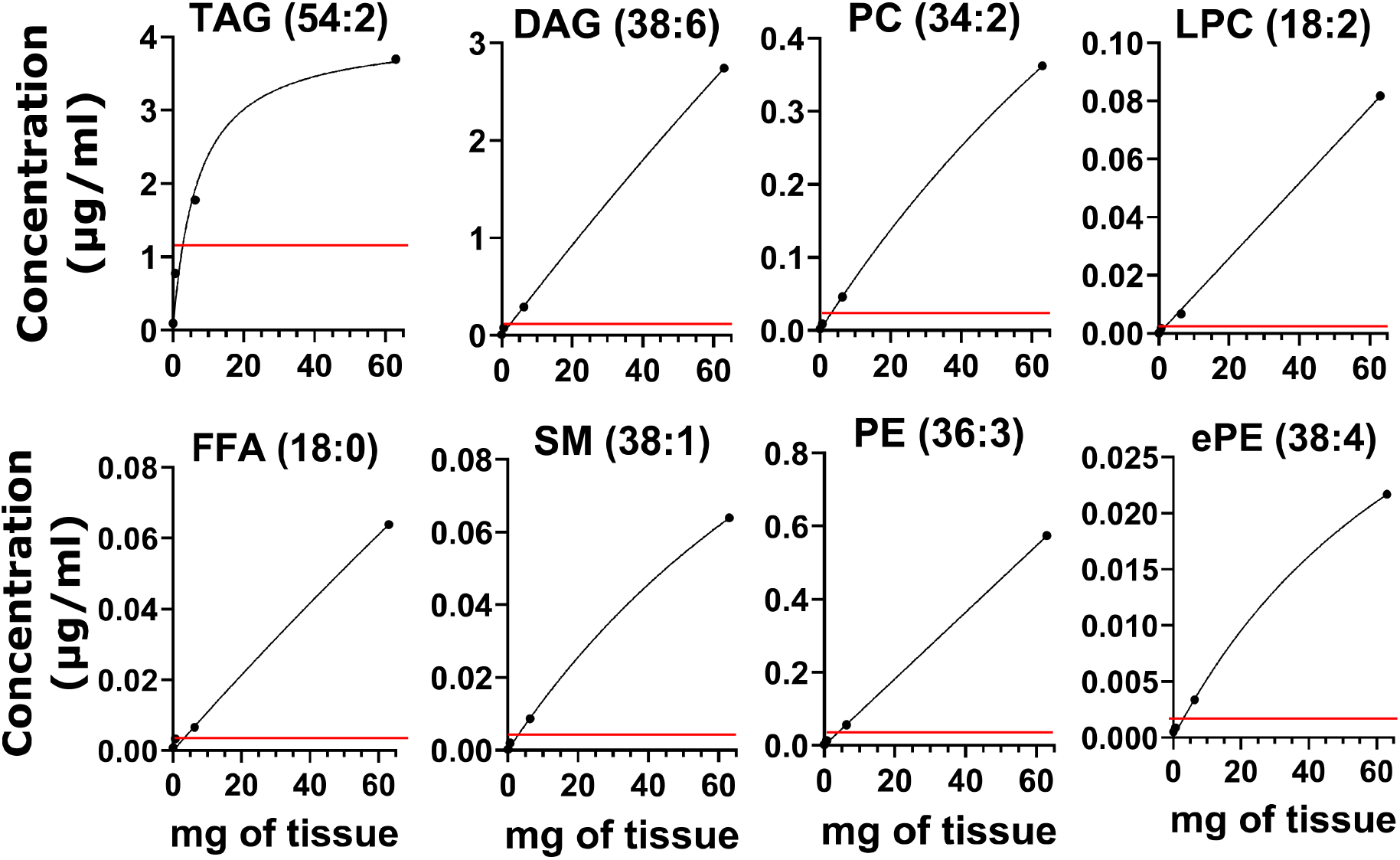
Linearity of various lipid groups by tissue biopsy weight. The more abundant lipid classes show excellent (R^2^ ≥ 0.99) linearity over a high dynamic range (∼0.5 – 50 mg), however, as triacylglycerides (TAGs) are the most abundant group present in WAT, biopsies over 10 mg show reduced linearity.

**Supplementary Figure 2.**
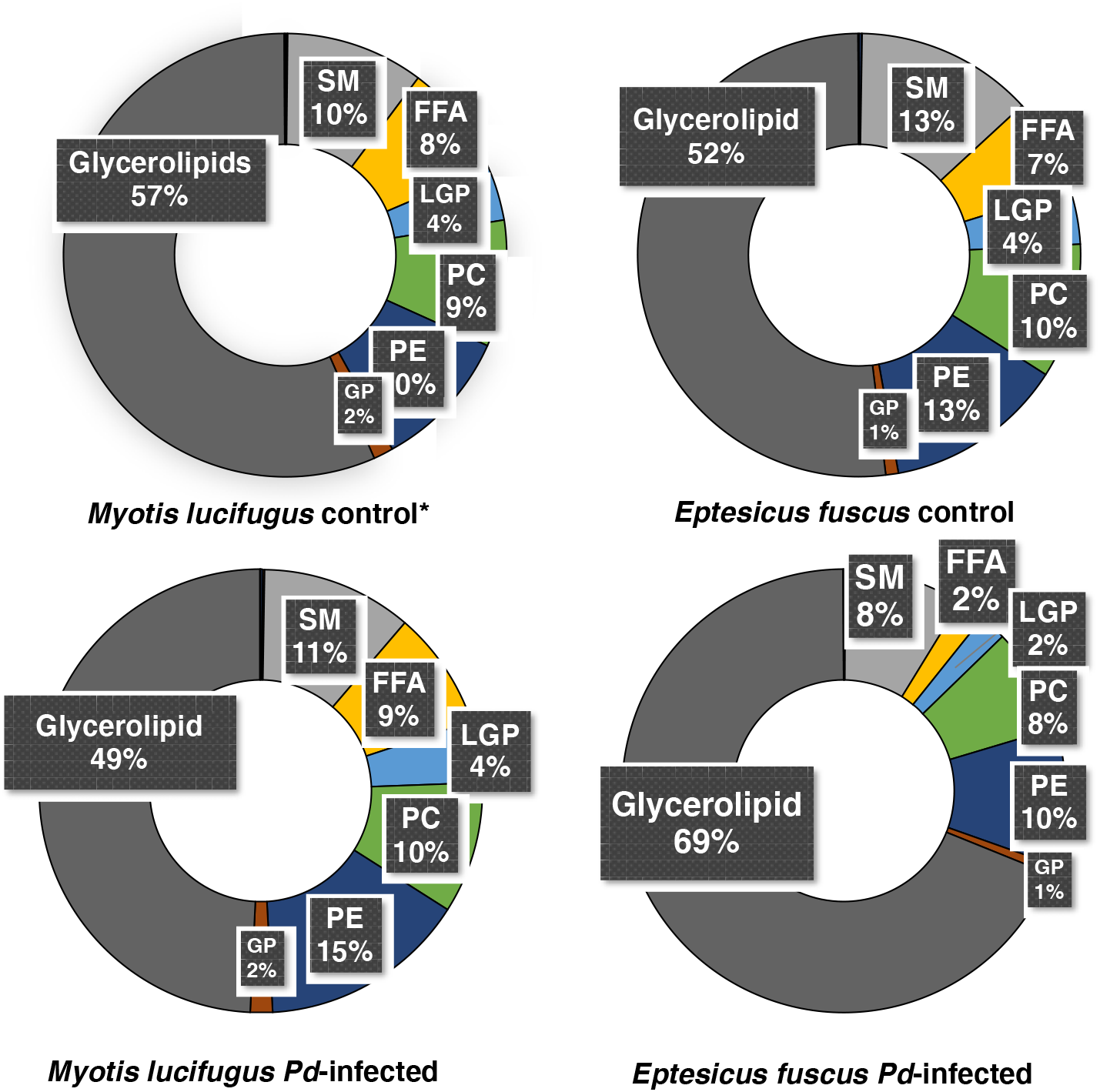
Donut plots depicting the chemical composition of white adipose tissue (WAT) in groups analyzed, including the control group for *Myotis lucifugus* (n=14)*, *Pd*-infected *M. lucifugus* (n=11), the control group for *Eptesicus fuscus* (n=6), and *Pd*-infected *E. fuscus* (n=5). * *M. lucifugus* control is from figure 1 and is provided here as a reference

**Supplementary Figure 3.**
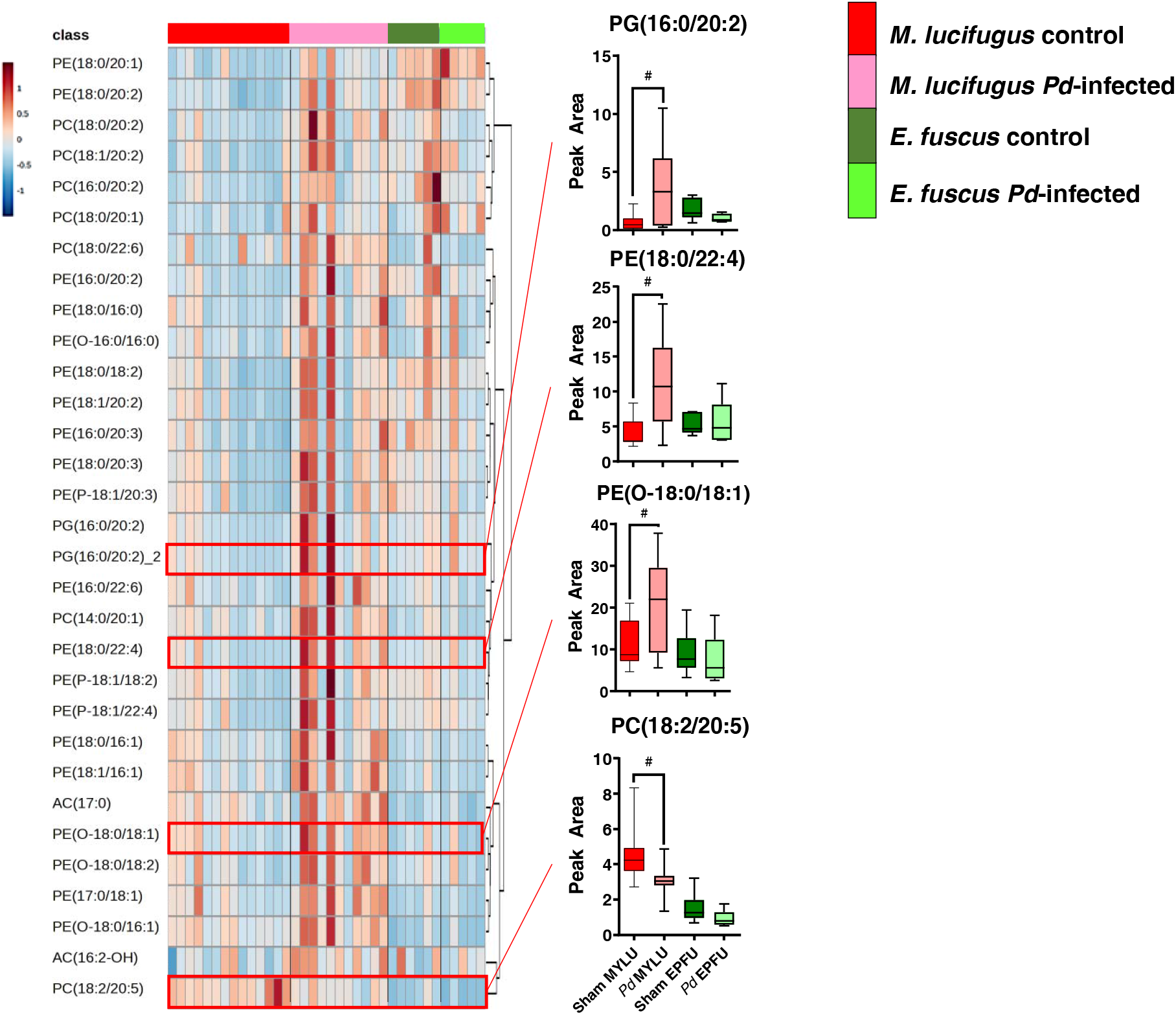
Individuals used to determine differences in lipid compounds in white adipose tissue (WAT) for Figure 4. (*FDR corrected P = 0.15, lines from top to bottom represent max value, 75th percentile, median, 25th percentile, min value)

